# Key role of vimentin in the organization of the primary cilium

**DOI:** 10.1101/2024.01.17.576004

**Authors:** Vasiliki Lalioti, Diego Moneo-Corcuera, Dolores Pérez-Sala

## Abstract

We previously reported the presence of the intermediate filament vimentin at the primary cilium of lung cancer epithelial cells. In this study we further demonstrate that vimentin is intimately intertwined with acetylated tubulin at this structure. Interestingly, although vimentin can be detected along the whole length of the primary cilium, phospho-serine 56 vimentin is found particularly enriched at its basal region in A549 lung cancer cells. Vimentin appears to play a pivotal role in ciliogenesis, since its depletion in MEF or in A549 cells results in a lower proportion of cells displaying primary cilia and recognizable basal bodies. Furthermore, the detectable cilia in vimentin depleted cells are shorter. In addition, the centriolar structure appears disrupted in vimentin deficient cells, as indicated by an abnormal distribution of γ- and acetylated tubulin. Moreover, these cells display a defective organization of the pericentriolar material, characterized by a marked decrease in the levels of pericentrin and a diffuse distribution of Rab11. Taken together, our results show that vimentin is present at the primary cilium and suggest that it plays an important role in cilium structure and biogenesis, since its depletion leads to marked morphological defects and altered organization of key elements of this structure.

## Introduction

Intermediate filaments are cytoskeletal structures that form versatile arrangements which play structural, regulatory and integrating cellular roles. Cytoplasmic intermediate filaments can form extended networks that give support to the cell and resistance against various types of stress, connect and keep the position and homeostasis of organelles, and participate in mechanosensing and cell-cell communication [1, 2]. Vimentin, a type III intermediate filament protein, provides a remarkable example of the multifaceted roles of these cytoskeletal structures [3, 4]. Vimentin is expressed in cells of mesenchymal origin, as well as in cells that undergo epithelial-mesenchymal transition during tumorigenesis. Vimentin is normally present in cells forming a robust filament network extending throughout the cytoplasm. Vimentin filaments are highly dynamic and can undergo assembly and disassembly, severing and reannealing, and subunit exchange along their entire length [5]. This dynamic remodeling is controlled by a complex array of posttranslational modifications that act in concert depending on the cell state and needs [6–8]. Vimentin is intimately connected to the other main cytoskeletal networks, formed by actin and tubulin, and plays a key role in the coordination of cellular responses [2, 9–11]. In addition to the thoroughly studied filamentous form, vimentin can be present as non-filamentous structures, both inside cells and at the cell surface, as well as in the extracellular medium, where it can appear in “soluble” form or in extracellular vesicles [3]. At these locations, vimentin plays a variety of functions different from its intracellular roles [12]. At the cell surface, vimentin can act as a receptor for various ligands, including soluble CD44 and various glycated proteins, and as a coreceptor for pathogens [3, 13]. Conversely, extracellular vimentin can also act as a ligand for a variety of receptors, such as various lectins and IGF-1R (reviewed in [3]). Interestingly, we recently reported the presence of vimentin at the primary cilium of lung cancer epithelial cells [14]. However, the functional implications of vimentin at this location have not been elucidated.

Primary cilia constitute unique structures in the form of “antennas” that protrude from resting cells and play important roles in sensing the environment, signal transduction, cell-cell communication and responses to stress. The primary cilium contains a microtubule-based scaffold, called the axoneme, which is surrounded by a plasma membrane of specialized composition, highly enriched in receptors and ion channels [15] (see scheme in Fig. 1A). The axoneme emerges from the basal body, which is a modified mother centriole that is connected to the daughter centriole. The basal body is formed by nine microtubule triplets arranged in a cylindrical disposition. The microtubule scaffold evolves from the base to the tip of the cilium [16], and, as the axoneme extends, the triplets become doublets and the number of microtubules further decrease towards the tip [16]. Microtubules in the axoneme are highly enriched in acetylated tubulin [17].

**Figure 1.**
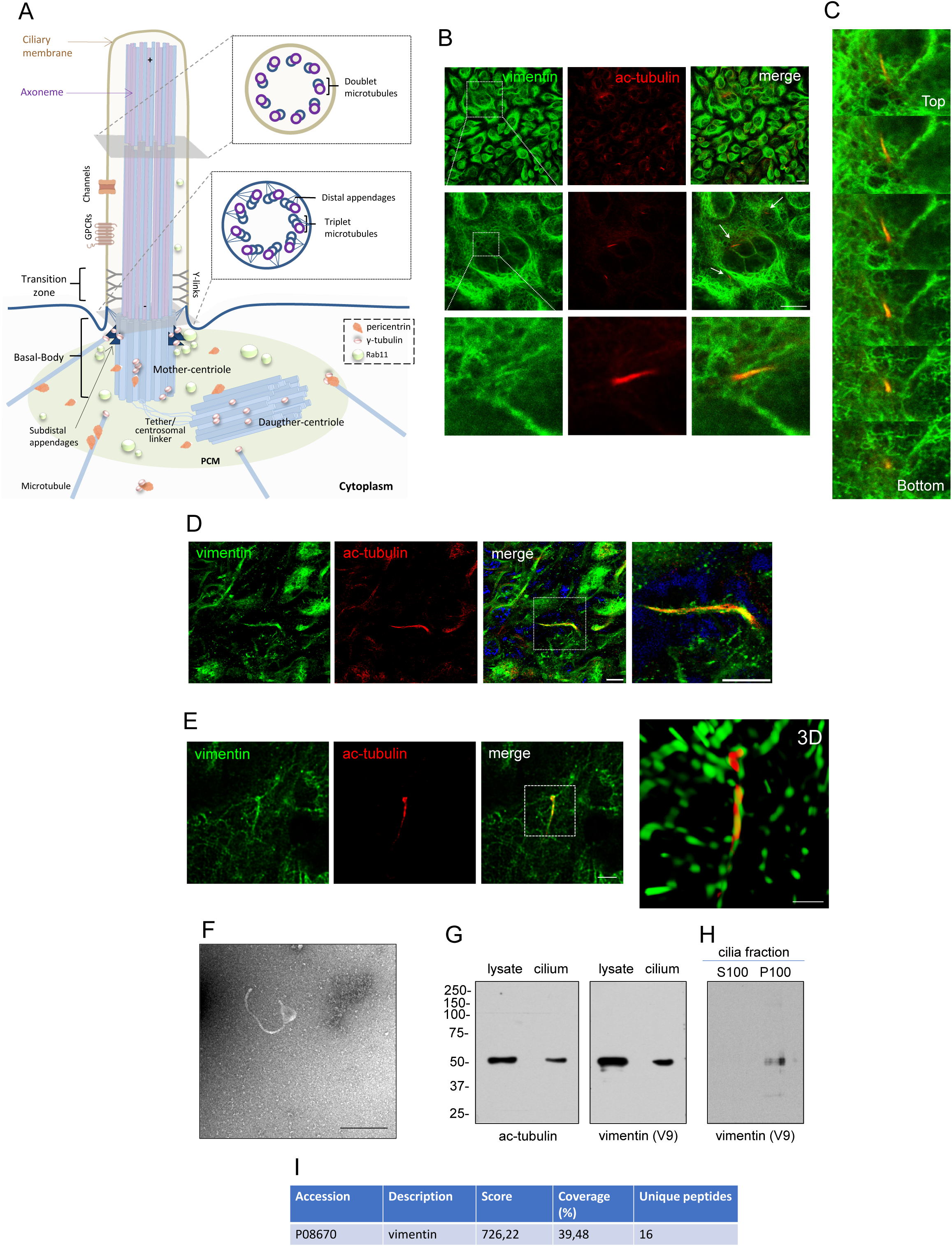
Presence of vimentin at the primary cilium. (A) Scheme of the architecture and main elements of the primary cilium. In quiescent cells the centrosome evolves to give rise to the primary cilium. The mother centriole forms the basal body from which nine microtubule triplets extend forming the internal cilium scaffold or axoneme. At distal sections the microtubule triplets become doublets. Tubulin is heavily acetylated along the length of the cilium. The two centrioles are surrounded by the pericentriolar material (PCM), which contains elements important for ciliogenesis, including Rab GTPases in charge of the traffic of building elements for the cilium, γ-tubulin and pericentrin (please see text for details). (B) A549 cells were cultured during 7 days after passage and vimentin (antibody 84.1) and acetylated tubulin (ac-tubulin) were detected by immunofluorescence. The upper row presents overall projections of the individual and merged channels. The area of interest is enlarged in the medium row, which depicts single confocal sections, where several structures compatible with cilia are marked by arrows. One of the cilia is shown in the lower row in more detail. Scale bars, 20 µm. (C) A stack of the single sections obtained every 0.5 µm is shown to illustrate the upward projection of the cilium. Images are rotated 90° counter-clock wise with respect to (B). (D) A549 cells were cultured and stained with the antibodies 84.1 and anti-acetylated tubulin, as above. Images were obtained with the Lightning module of the Leica SP8 microscope. Single channels and merged images are shown. The region of interest denoted by the dotted square is enlarged at the right (scale bars, 10 and 5 μm, respectively). (E) Images of a cilium were obtained by STED superresolution microscopy and a 3D reconstruction of this structure is provided in the right image to illustrate the close intertwining of vimentin and acetylated tubulin (scale bars, 5 and 2 μm, respectively). (F) The cilia fraction was isolated and analyzed by electron microscopy; scale bar, 500 nm. (G) Total lysates from A549 cells and the cilia fraction were analyzed by SDS-PAGE and western blot with the indicated antibodies. (H) The cilia fraction was resuspended in hypotonic buffer and soluble (S100) and insoluble (P100) fractions, analyzed by SDS-PAGE and western blot with anti-vimentin. (I) The cilia fraction was subjected to proteomic analysis and the parameters corresponding to the identification of vimentin are shown.

The process of ciliogenesis is highly complex. When cells exit the cell cycle, the mother centriole of the centrosome undergoes a remodeling process to become the basal body. Ciliogenesis can occur through different routes, intra or extracellular, depending on the cell type [18, 19]. In the intracellular pathway, vesicles containing Rab 11 and Rab 8 GTPases converge at the distal end of the mother centriole to form a ciliary vesicle [20, 21]. The capping proteins of the mother centriole are removed by proteasomal degradation and this allows extension of the microtubules that grow pushing this vesicle outwards and causing the cilium to protrude from the plasma membrane [19, 22]. At the same time, the membrane remodels to wrap the axoneme and becomes the ciliary membrane. The base of the cilium becomes a very complex structure that contains the basal body, with appendages that dock it to the membrane, and a region known as the pericentriolar material [23]. Besides the basal body, the pericentriolar material enfolds multiple elements, including the daughter centriole, proteins of the Rab family of GTPases, which regulate the vesicular transport of materials required for cilium growth to this area, pericentrin, a centrosomal scaffold protein that contributes to organize microtubules, and γ-tubulin complexes [24]. Both the basal body and the daughter centriole contain γ-tubulin ring complexes (γ-TuRC) [25]. In addition, multiple microtubules radiate from this location, which are nucleated by γ-TuRC and decorated by pericentrin [25] (Fig. 1A).

Although the various forms of tubulin are essential architectural and regulatory components of the primary cillium, actin also plays an important role in the regulation of cilium properties [26, 27]. In addition, other cytoskeletal elements, including septins and intermediate filament-associated proteins, have been reported to be present at the primary cilium and/or to influence its formation [27–31]. The results herein described confirm our previous observations on the presence of vimentin at diverse structures of the primary cilium [14], and indicate that vimentin is required for ciliogenesis, since blocking vimentin expression leads to severe defects in cilia morphology and in the organization of key cilia structural and regulatory components. Nevertheless, this picture could be more complex, given the sophisticated interplay between intermediate filaments and the actin and tubulin cytoskeletal structures.

### Experimental section

#### Reagents

Buffers, paraformaldehyde (PFA), triton X-100, and methanol were from Merck. 4’, 6-Diamidino-2-phenylindol (DAPI) was from Sigma. A full description of the antibodies used is provided as Suppl. Table 1.

#### Cell culture

Lung cancer adenocarcinoma A549 cells (ATCC, CCL-185) were cultured in RPMI1640 with 10% (v/v) FBS, 50 U/ml penicillin, 50 μg/ml streptomycin and 50 μg/ml gentamycin. Cells were additionally authenticated by microsatellite sequencing at Secugen (Madrid, Spain). A549 *VIM*KO cells were obtained by disruption of the *VIM* gene with the CRISPR double nickase technology using the Vimentin Double Nickase Plasmid (h) from Santa Cruz Biotechnology (sc-400035-NIC), following the instructions of the manufacturer. Mouse embryonic fibroblasts (MEF) from wild type (MEF wt) and vimentin knockout mice (MEF *Vim(-/-)*) were the generous gift of Prof. John Eriksson (Abo Academy, Turku, Finland). Cells were cultured in DMEM supplemented with 10% (v/v) fetal bovine serum (FBS, Sigma) and 100 U/ml penicillin, 100 μg/ml streptomycin. To obtain quiescent cells with developed cilia, unless stated otherwise, cells were cultured for five days after plating and then subjected to partial serum starvation in medium containing 0.5% (v/v) FBS for two more days in the case of A549 cells or overnight for MEF. All cell lines were periodically confirmed to be free of mycoplasma contamination at the Animal Cell Culture facility of the CIB Margarita Salas (Madrid, Spain).

#### Plasmids and transfections

The GFP-vimentin wt fusion plasmid has been already reported [32, 33]. Cells were transfected with Lipofectamine 2000 (Invitrogen), following the instructions of the manufacturer. Typically, 0.2-0.5 µg of DNA for A549 cells and 1 µg of DNA for MEF plus 3 µl of Lipofectamine were used for transfection of cells in a p35 dish.

#### Isolation of cilia fraction

Cilia were obtained essentially as described [34]. Cells were seeded on p100 dishes and cultured as above for cilia development. Cell culture medium was carefully removed and cells were gently washed with cold PBS. Subsequently, cells were washed with 10 ml of PBS by rotary shaking at 360 rpm for 5 minutes to detach cilia. This medium was collected and centrifuged at 1000 g for 10 min at 4°C to remove cell debris. The supernatant was centrifuged at 40000 g for 30 min at 4°C, and the pellet containing cilia was resuspended in 20 mM Tris HCl pH 8, 50 mM KCl, 4 mM MgSO4, 1 mM DTT, 0,5 mM EDTA for analysis by electron microscopy or SDS-PAGE, followed by immunodetection or proteomic analysis. When indicated, cilia were resuspended in hypotonic buffer, 5 mM Pipes pH 7.0, for further analysis.

#### Electron microscopy

Briefly, aliquots of the cilia fraction were fixed by addition of 0.1% (v/v) glutaraldehyde, final concentration. Then, 5 µl aliquots of cilia fraction were adsorbed onto carbon support grids (MESH CF 400 CU UL, Aname), which were subsequently washed with water and stained with 2% (w/v) uranyl acetate. Grids were inspected on a JEOL transmission electron microscope JEM-1230, equipped with a digital camera CMOS TVIPS TemCam-F416, at the Electron Microscopy facility of CIB Margarita Salas.

#### Immunofluorescence

Cells grown on glass coverslips were fixed by incubation with 4% (w/v) paraformaldehyde (PFA) for 10 min at room temperature, and permeabilized with 0.2% (w/v) triton X-100 for 5 min. Following these steps, reactive aldehydes were quenched by incubation with 10 mM glycine for 20 min. After blocking with 1% (w/v) BSA in PBS, cells were incubated with the primary antibodies of interest, usually at 1:300 dilution in blocking solution, followed by secondary antibodies conjugated with Alexa647 or 488, at 1:200 dilution, essentially as described [14]. Detection of vimentin was achieved by direct or indirect immunofluorescence, depending on the antibody used, which included clones 84.1, SP20, V9-A488, and E5-488. Detection of acetylated tubulin was achieved by incubation with anti-acetylated tubulin-A546 at 1:400 dilution. In all cases, controls without primary or secondary antibodies were processed in parallel to ensure the specificity of the signals. Nuclei were counterstained with DAPI at 2.5 μg/ml in PBS. Washing steps were performed by careful immersion of coverslips in PBS in order not to disrupt cilia. At the end of the procedure coverslips were mounted with Fluorsave (Millipore).

#### Confocal microscopy and image analysis

Cells on glass coverslips were examined on Leica SP5 or SP8 confocal microscopes. Routinely, sections were acquired every 0.5 µm in sequential mode, and overall projections or single sections are shown, as indicated. For higher resolution, either the Lightning module of the SP8 microscope with Adaptive setting or STED superresolution microscopy were employed. The LUT command was used to ensure acquisition under non-saturated conditions. Nevertheless, when indicated, images provided are deliberately overexposed to illustrate specific features. Image J was used for quantitation. For assessment of the number and length of cilia, well-delimited elongated structures positive for acetylated tubulin were considered.

#### Cell lysis and western blot

Cells were washed twice with cold PBS before lysis in a RIPA buffer containing cOmplete™ Protease Inhibitor Cocktail (Sigma). Cell debris was removed by centrifugation at 16000 g for 5 min at 4°C. Protein concentration was determined by the Bicinchoninic acid method (Pierce, ThermoFisher Scientific). Aliquots of lysates containing 30 μg of protein were separated on SDS-PAGE and transferred to Immobilon-P membranes (Millipore) using a Trans-Blot semi-dry transfer unit from Bio-Rad and a three buffer sandwich, as indicated by the manufacturer. Before immunodetection, blots were blocked with 2% (w/v) non-fat evaporated milk in T-TBS. Primary antibodies were typically used at 1:500 dilution, followed by horseradish peroxidase conjugated secondary antibodies, at 1:2000 dilution. Bands of interest were visualized with the enhanced chemiluminiscence system (ECL, GE Healthcare), by exposure to Agfa films.

#### LC-MS analysis

Peptide separations were carried out on an Easy-nLC 1000 nano system (Thermo Scientific). For the analysis, the sample was loaded into a precolumn Acclaim PepMap 100 (Thermo Scientific) and eluted in a RSLC PepMap C18, 50 cm long, 75 µm inner diameter and 2 µm particle size (Thermo Scientific). The mobile phase flow rate was 300 nl/min using 0.1% formic acid in water (solvent A) and 0.1% formic acid and 100% acetonitrile (solvent B). The gradient profile was set as follows: 5–35% solvent B for 100 min, 35%-45% solvent B for 20 min, 45%-100% solvent B for 5 min, and 100% solvent B for 15 min. Four microliters (1 µg) of each sample were injected. MS analysis was performed using a Q-Exactive mass spectrometer (Thermo Scientific). For ionization, 1900 V of liquid junction voltage and 250 °C capillary temperature was used. The full scan method employed a m/z 300–1800 mass selection, an Orbitrap resolution of 70,000 (at m/z 200), a target automatic gain control (AGC) value of 3e^6^, and maximum injection times of 100 ms. After the survey scan, the 15 most intense precursor ions were selected for MS/MS fragmentation. Fragmentation was performed with a normalized collision energy of 27 eV and MS/MS scans were acquired with a starting mass of m/z 200, AGC target was 2e^5^, resolution of 17500 (at m/z 200), intensity threshold of 8e^3^, isolation window of 2.0 m/z units and maximum IT was 100 ms. Charge state screening was enabled to reject unassigned, singly charged, and equal or more than seven protonated ions. A dynamic exclusion time of 20s was used to discriminate against previously selected ions.

#### MS data analysis

MS data were analyzed with Proteome Discoverer (version 1.4.1.14) (Thermo) using standardized workflows. Mass spectra *.raw files were searched against the SwissProt 2021_01, Taxonomy Homo sapiens (20397 sequences) using Mascot search engine (version 2.6, Matrix Science). Precursor and fragment mass tolerance were set to 10 ppm and 0.02 Da, respectively, allowing 2 missed cleavages, carbamidomethylation of cysteines as a fixed modification, serine, threonine, tyrosine phosphorylation methionine oxidation, and acetylation N-terminal as a variable modification. Identified peptides were filtered using Percolator algorithm [35] with a q-value threshold of 0.01. The protein identification by nLC-MS/MS was carried out in the Proteomics and Genomics Facility (CIB-CSIC), a member of ProteoRed-ISCIII network.

#### Statistical analysis

All experiments were repeated at least three times. Statistical analysis was performed with GraphPad version 9, and differences between experimental conditions were assessed by t-test or ANOVA.

## Results

### Presence of vimentin at primary cilia

The primary cilium is a complex structure organized around the axoneme. A schematic view of the primary cilium is shown in Fig. 1A. We previously showed that vimentin could be immunodetected at the primary cilium of A549 lung cancer cells, where it colocalized with well-known cilia markers such as acetylated tubulin and the small GTPase ARL13b [14]. Here we have confirmed the presence of vimentin at these structures through several approaches. Immunofluorescence of vimentin in A549 cells clearly detects its presence at typical primary cilia appearing as elongated structures, where it associates with acetylated tubulin (Fig. 1B). These structures protrude from cells, as illustrated in Fig. 1C, which depicts a stack of single sections, from the putative position of the basal body (bottom image) to nearly the tip of the cilium (top image). The association of vimentin and acetylated tubulin along the length of the cilium can be appreciated. Vimentin localization at the primary cilium can also be observed in A549 cells transfected with GFP-vimentin (Suppl. Fig. 1A), although it was more obvious in cells with lower GFP-vimentin levels. The intimate association of acetylated tubulin and vimentin can be further evidenced by employing the Lightning module of the SP8 confocal microscope (Fig. 1D). Interestingly, in some cases, vimentin appears to wrap around the acetylated tubulin signal, as revealed by STED superresolution microscopy (Fig. 1E). Indeed, 3D reconstruction of these images clearly shows that vimentin structures extend throughout the length of the cilium intertwining with acetylated tubulin (Fig. 1E, right image). Finally, as cilia are flexible structures it can be difficult to capture their whole length in a single confocal section; therefore, it is important to note that the apparent fading of the vimentin and acetylated tubulin signals at certain points along the length of the cilium is likely due to their presence in a different confocal plane (Suppl. Fig. 1B).

Electron microscopy of the cilia fraction isolated from A549 cells by gentle shaking shows the presence of elongated structures compatible with primary cilia [34] (Fig. 1F). Moreover, the results from western blot confirm the presence of vimentin and acetylated tubulin in the analyzed cilia fraction (Fig. 1G). The nature of the vimentin structures present in this fraction was further explored by resuspension in hypotonic buffer and ultracentrifugation. This showed that vimentin was only detected in the pellet, indicating its presence in insoluble oligomeric or polymeric forms (Fig. 1H). Finally, the cilia fraction was subjected to proteomic analysis, which confirmed the presence of vimentin. The parameters corresponding to the identification of vimentin are summarized in Fig. 1I, and the sequence of the peptides identified is provided in Suppl. Table 2. In addition to vimentin, other proteins were identified in this fraction, including plectin, filamin, actin, tubulin, several Rab GTPases, ion channels and receptors, as well as several chaperones, the presence of which in intact cilia in cells requires further confirmation. The complete list of proteins identified is provided in Suppl. Table 3.

Collectively, these results confirm the presence of vimentin within the primary cilia of A549 cells, revealing a close intertwining with acetylated tubulin.

### pSer56-vimentin is selectively associated with primary cilia

Vimentin organization and subcellular distribution are tightly regulated by phosphorylation [6, 36]. In turn, vimentin phosphorylation at certain residues can occur at specific subcellular locations [37]. Therefore, we employed three different anti-phosphovimentin antibodies to explore the distribution of phosphorylated vimentin in relation to the primary cilium in A549 cells (Fig. 2). The numbers of the phosphoresidues refer to the sequence of human vimentin including the initial methionine. We observed that the three antibodies used yielded different patterns (Fig. 2A). Anti-pSer39-vimentin yielded a mostly filamentous pattern which markedly coincided with the total vimentin network stained with the SP20 anti-vimentin antibody. In contrast, anti-pSer72 and anti-pSer56-vimentin displayed a more discontinuous staining. In particular, the anti-pSer56-vimentin antibody recognized numerous punctate accumulations and elongated structures which could correspond to cilia. Indeed, co-staining with acetylated tubulin clearly showed that the pSer56-vimentin antibody highlighted primary cilia and/or regions apparently located near its basal region (Fig. 2B). This selective localization was confirmed in MEF wt, although in this cell type, staining of cilia with anti-pSer56-vimentin appeared to be fainter and less intense at the basal region (Suppl. Fig. 2A). Of note, we also observed an enrichment of the pSer56-vimentin signal throughout the vimentin network in cells undergoing mitosis, either MEF or A549 cells (Suppl Fig. 2A and B). This is consistent with the phosphorylation of this site by CDK1, reported earlier in mitotic cells [38]. Therefore, pSer56-vimentin appears to highlight different structures in resting and mitotic cells.

**Figure 2.**
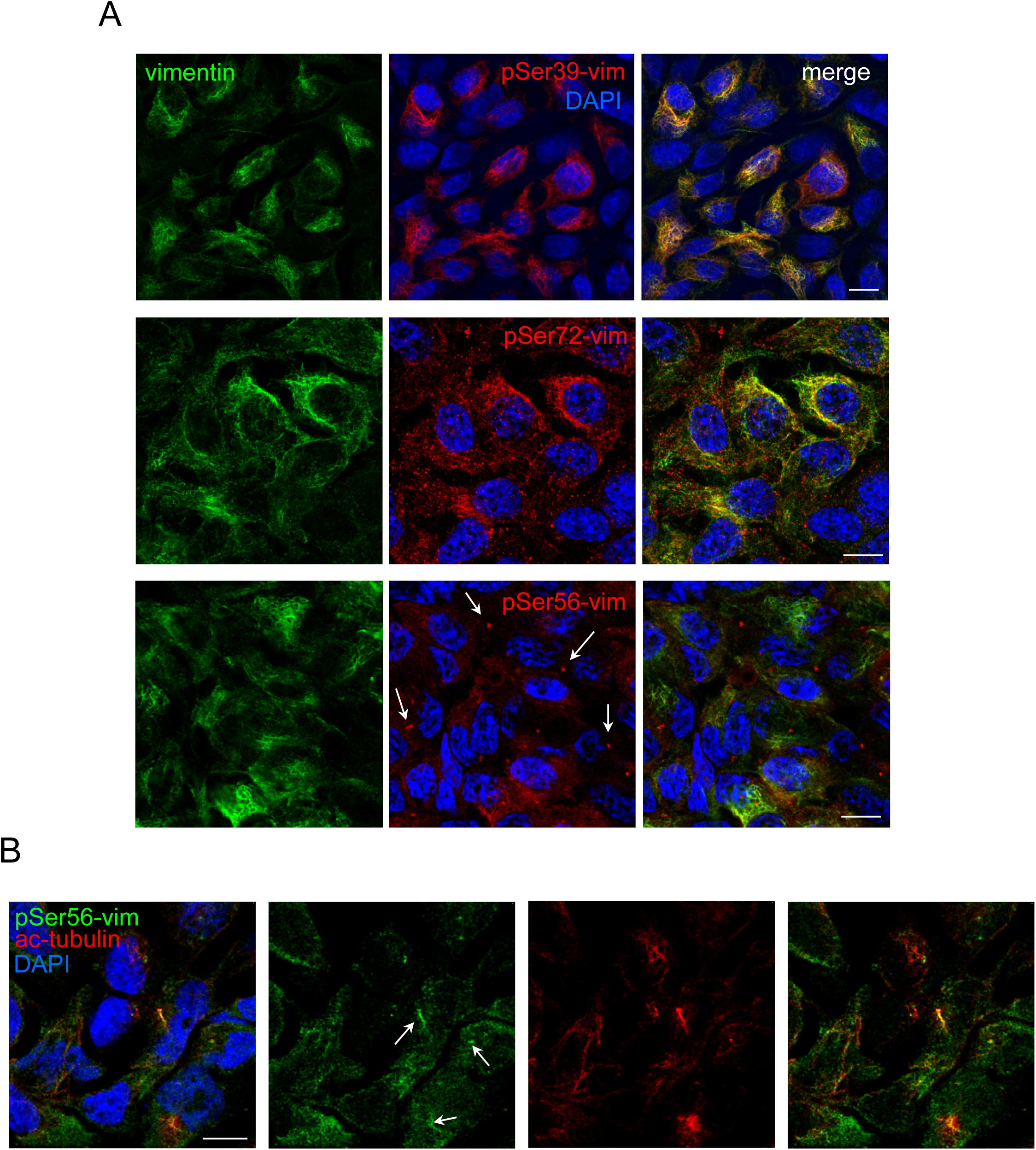
Detection of phosphorylated vimentin in association with primary cilia. (A) A549 cells cultured as above were fixed and processed for immunofluorescence. Total vimentin was detected with the SP20 antibody and phospho-specific antibodies were used for detection of pSer39, pSer56 or pSer72-vimentin. In the lower row, signals compatible with primary cilia are marked by arrows. (B) Localization of pSer56-vimentin at primary cilia was confirmed by staining of acetylated tubulin. Images shown are single confocal sections. Several structures positive for both proteins are marked by arrows. Bars, 20 μm.

In view of these results, we next explored the specific localization patterns of pSer56-vimentin at the primary cilium, seeking a more comprehensive understanding (Fig. 3A). We observed a marked colocalization of robust pSer56-vimentin and acetylated tubulin signals along the cilia length in A549 cells. Interestingly, a strong pSer56-vimentin signal was also observed at the basal region of the cilia, coinciding with a weak acetylated tubulin staining. Further magnification and exposure of these images illustrated that the basal and ciliary signals were connected. This suggested that pSer56-vimentin was enriched at the origin of the cilia. To confirm this possibility, we explored its colocalization with Rab11, which is known to be enriched at the base of primary cilia [20]. In A549 cells, Rab11 yielded a punctate pattern distributed throughout the cytoplasm, but displayed a defined accumulation coinciding with the base of primary cilia, which were detected with anti-acetylated tubulin antibody (Fig. 3B). A higher magnification showed the disposition of Rab11 in “U” or ring-like shaped formations encircling the proximal region of the acetylated tubulin signal (Fig. 3B). Interestingly, this Rab11 enriched region appeared in close proximity, and at some points surrounding the pSer56-vimentin signal (Fig. 3C), suggesting that pSer56-vimentin is accumulated at or in the proximity of the basal body or in the pericentriolar area. Additional images obtained with the Lightning module for higher resolution, which illustrate the disposition of Rab11 around pSer56-vimentin in the basal region of the cilia are shown in Suppl. Fig. 3. Interestingly, antibodies against total vimentin highlighted mainly the region of the axoneme, yielding a less distinctive signal at the basal region, which partially colocalized with the Rab11-positive signal (Fig. 3D). Taken together, these observations suggest that selective phosphovimentin proteoforms, in particular pSer56-vimentin, appear to be enriched at primary cilia, and more precisely, at the basal region, in comparison with total vimentin.

**Figure 3.**
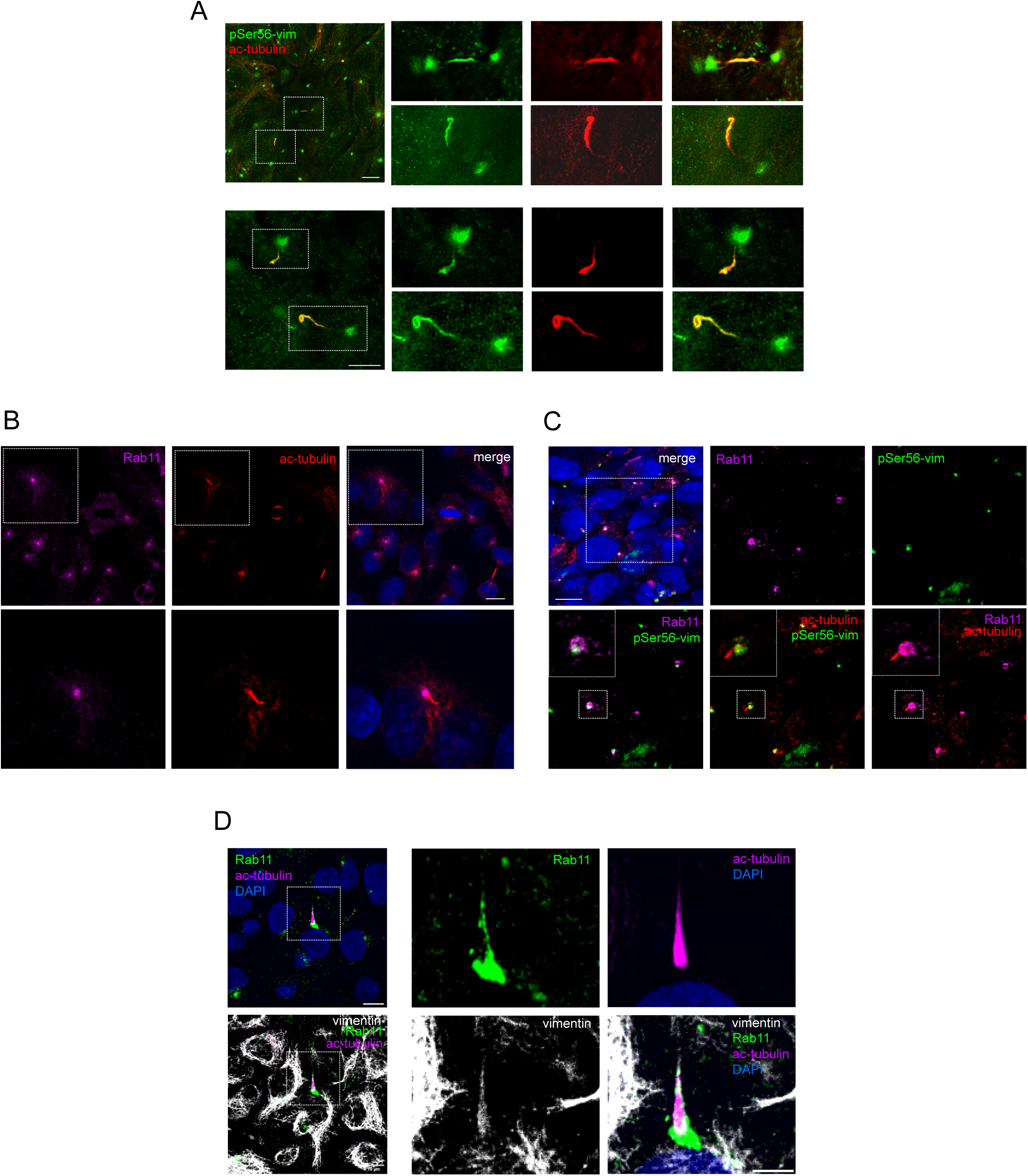
Characterization of the presence of pSer56-vimentin at the primary cilium. (A) A549 cells were cultured for an initial 5-day period and then subjected to a starvation condition for two days. pSer56-vimentin and acetylated tubulin were detected by immunofluorescence and STED microscopy. Areas of interest denoted by dotted boxes are enlarged at the right (scale bars, upper image 10 µm, lower image, 5 µm). (B) Immunostaining of Rab11 and acetylated tubulin to show the presence of Rab11 at the base of the primary cilium. Inset is enlarged at the bottom images. Images shown are single sections. (C) The presence of the pSer56-vimentin signal at the pericentriolar material is assessed by immunofluorescence using Rab11 as a marker of this structure. The area of interest in the merged image is enlarged at right and at the bottom. Further detail of one of the cilia is depicted in insets. Images shown are single sections. (D) Detection of total vimentin in relation to the pericentriolar material. Images at the right depict composite images of one cilium stained with anti-Rab11 (green), anti-acetylated tubulin (magenta), and for total vimentin (84.1 antibody, grayscale). Nuclei were counterstained with DAPI. Images at the right show enlarged views of the area of interest as single channels for the various signals and a merged image (bottom right). Note the presence of vimentin at the cilium and the absence of enrichment of its signal at the pericentriolar material region highlighted by the Rab11 signal. Images shown are single sections. Scale bars, left images, 10 µm, right images, 5 µm.

### Role of vimentin in cilia morphology

In order to explore a potential functional role of vimentin at the primary cilium, we analyzed the morphology of this organelle in vimentin expressing and vimentin depleted cells (Fig. 4). The lack of detectable vimentin in MEF *Vim(-/-)* was confirmed by immunofluorescence, as shown in Fig. 4A (lower images). MEF expressing vimentin were characterized by the presence of cilia with regular morphology in a high proportion of cells, as visualized by acetylated tubulin staining, which showed a certain degree of overlap with the vimentin signal. In sharp contrast, in the case of MEF from vimentin knockout mice (MEF *Vim(-/-)*), the proportion of cells showing well-formed cilia was significantly lower (Fig. 4A, graph). Moreover, the cilia that could be detected in MEF *Vim(-/-)* displayed irregular morphology and shorter length (Fig. 4B and graph). In particular, acetylated tubulin yielded a disorganized pattern, characterized by a higher background, together with fibers frequently stemming from the expected position of the basal body, as illustrated in Fig. 4B, lower images.

**Figure 4.**
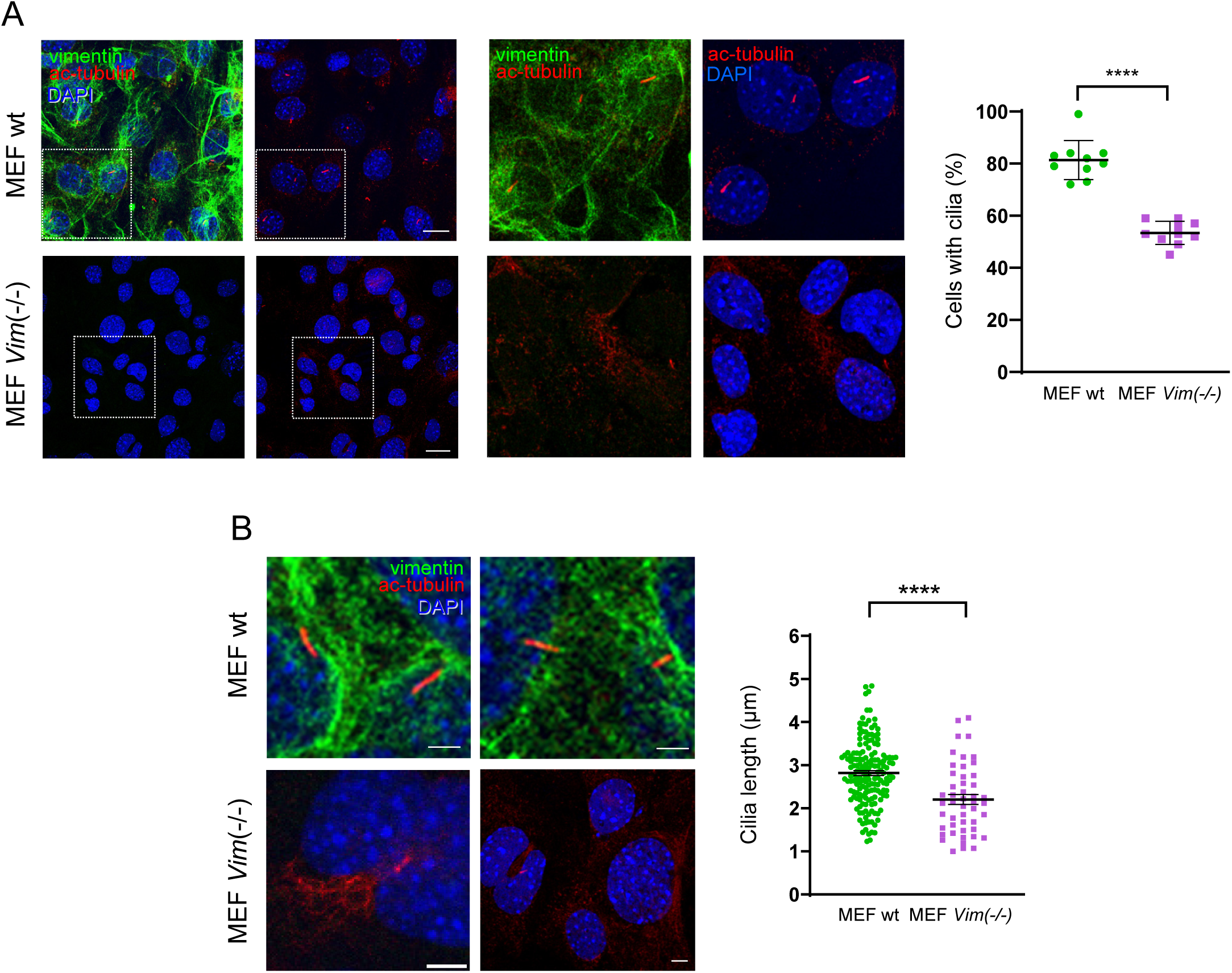
Impact of vimentin knockout on the number and length of cilia. (A) MEF wt or *Vim(-/-)* were stained for total vimentin (E5 antibody) and acetylated tubulin. Areas of interest delimited by dotted boxes are enlarged in the images at the right. Images shown are single sections. Scale bars, 20 µm. The proportion of cells with detectable cilia in every cell type is shown in the graph at the right. Ten fields per experimental condition with 50 cells each were analyzed. Results shown are average values ± SEM of the 10 determinations. (B) Additional images illustrating the morphology of cilia encountered in MEF wt or *Vim(-/-)* immunostained as in (A). Note the presence of abundant acetylated tubulin fibers in the region corresponding to the base of cilia in MEF *Vim(-/-)*. Scale bars, 5 µm. The length of the detectable cilia, i.e., defined elongated structures (length ≥ 1 µm) positive for acetylated tubulin, in every cell type is quantitated in the graph at the right. Punctate or fibrous structures appearing in MEF *Vim(-/-)* were not considered. Results shown are determinations from three independent experiments, totaling 164 cilia for MEF wt and 85 for MEF *Vim(-/-)*. ****p<0.0001 by unpaired *t*-test.

These observations suggested a defect in ciliogenesis in vimentin knockout cells. Therefore, we evaluated if vimentin deficiency affected the organization of various primary cilia structures. The centrosomal protein γ-tubulin is an essential element in nucleation of microtubules. It forms the γ-tubulin ring complex, also known as γ-tuRC, a ring-like structure which serves to nucleate microtubules, both at the centrosome in interphase cells, and at the basal body of the primary cilium when cells exit the cell cycle. In addition, γ-tuRC can modulate cilia disassembly [19, 39]. In MEF wt γ-tubulin showed a clearly defined localization, frequently appearing as two adjacent spots at the base of the cilium, typical of its presence in centrioles [25] (Fig. 5A). In drastic contrast, MEF *Vim(-/-)* displayed an irregular and abnormal γ-tubulin distribution (Fig. 5A), which appeared as spots not related to the cilia, or as multiple spots located near the base of the abnormal cilia, and in some cases, even as an irregularly shaped signal entangled with the disorganized acetylated tubulin positive structures, as illustrated in the additional examples shown in Fig. 5B. These observations suggest the presence of severe centriole alterations in vimentin deficient cells.

**Figure 5.**
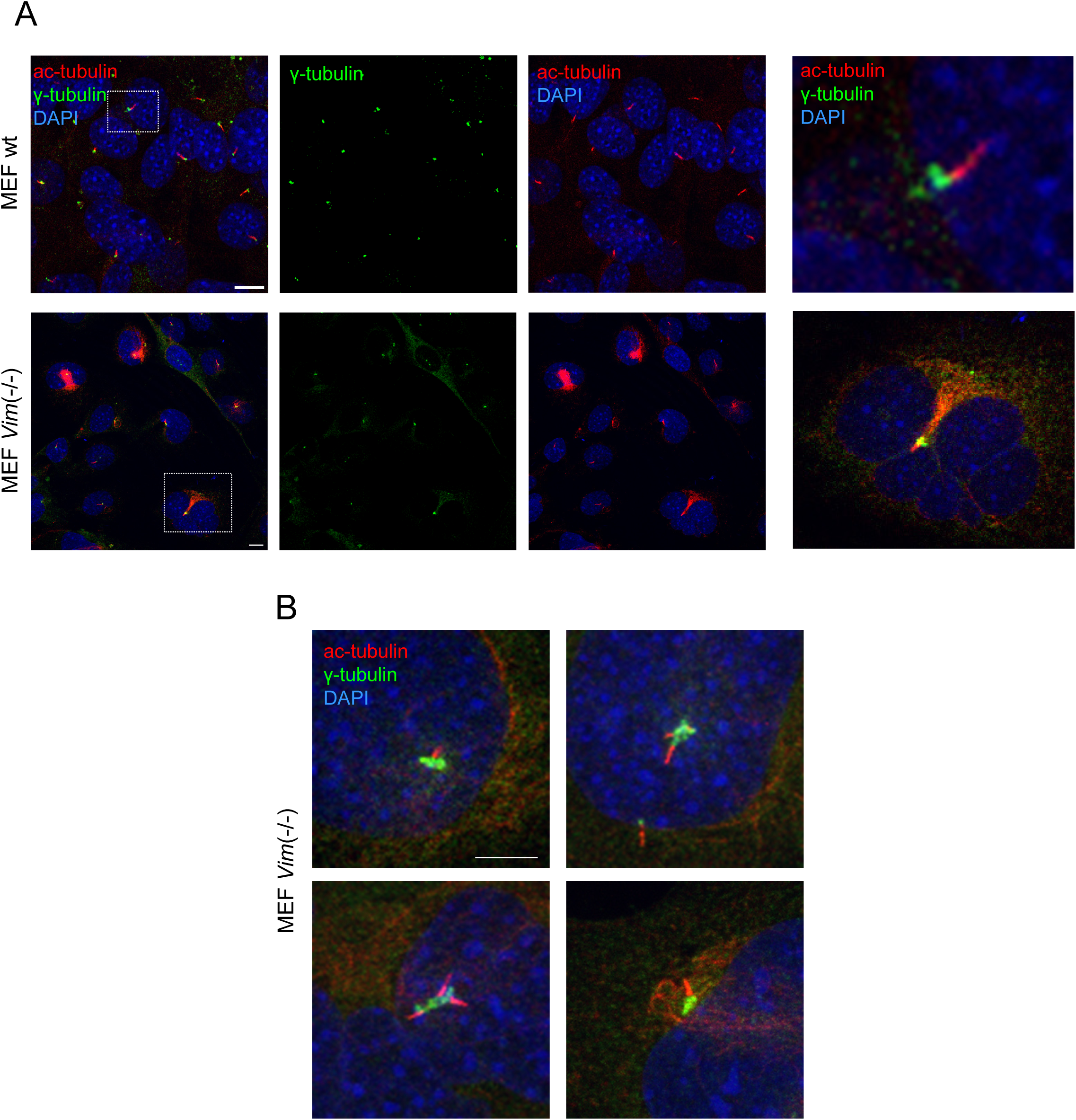
Impact of vimentin knockout on the organization of the centrioles. (A) MEF wt or *Vim(-/-)* were stained for γ-tubulin to highlight the centrioles and acetylated tubulin to locate the cilia. Single channels or overlay images are shown as indicated. The areas in dotted boxes are enlarged in the far-right images. Images shown are overall projections. Scale bars, 20 µm. (B) Additional images illustrating the altered morphology of centrioles in MEF *Vim(-/-)*. Note the disorganized appearance of γ-tubulin and acetylated tubulin-positive structures in these cells. Scale bars, 5 µm.

### Role of vimentin in pericentrin localization

In view of the abnormal distribution of both acetylated tubulin and γ-tubulin observed in MEF *Vim(-/-)*, we explored the distribution of pericentrin, a protein critical for the formation of primary cilia, which controls the traffic of intracellular material around the centrosome [40], and contributes to nucleate the microtubules by interacting with γ-tubulin [41]. Pericentrin is normally located around the centrioles at the base of primary cilia and its depletion impairs ciliogenesis [41]. Here we observed that in MEF wt pericentrin displayed a defined pattern, colocalizing with acetylated tubulin at the base of well developed cilia in a high proportion of cells (Fig. 6). In contrast, this distribution was profoundly altered in MEF *Vim(-/-)*, in which the pericentrin signal was significantly weaker (Fig. 6, graph) and appeared in less defined structures. Moreover, although in some MEF *Vim(-/-)* pericentrin appeared associated with acetylated tubulin, a well formed cilium could not be detected in most of them, as described above.

**Figure 6.**
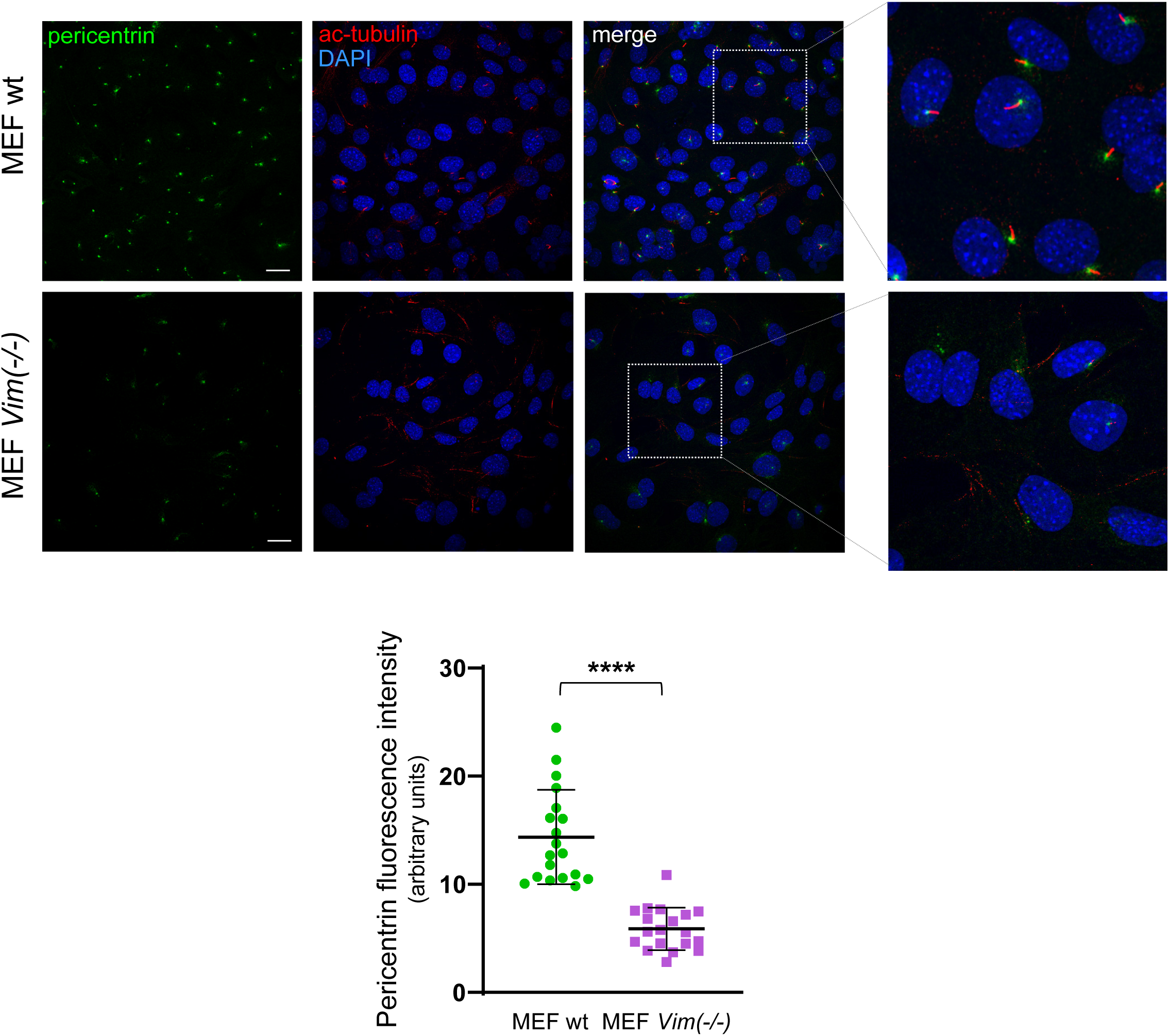
Impact of vimentin knockout on the distribution of pericentrin. (A) MEF wt or *Vim(-/-)* were stained for pericentrin and acetylated tubulin to locate the base of the cilia and the axoneme, respectively. Single channels or overlay images are shown as indicated. The areas in dotted boxes are enlarged in the far-right images. Images shown are overall projections. Scale bars, 20 µm. The intensity of the pericentrin fluorescent signal in MEF wt or *Vim(-/-)* is quantitated in the graph below. Twenty images with a similar number of nuclei were selected for every experimental condition from at least three independent experiments, and the intensity of the pericentrin signal was measured. Results are average values ± SEM. ****p<0.0001 by unpaired *t*-test.

### Vimentin depletion leads to defects in Rab11 localization at the pericentriolar material

Rab11 has been shown to play a key role in ciliogenesis [20], where it mediates the recruitment of elements needed for cilia assembly. Indeed, Rab11 has been reported to be required for pericentrin to be enriched at the centrosome [42, 43]. As we have described above (Fig. 3), a proportion of cellular Rab11 concentrates at the basal region of the cilia in wt cells. In view of the profound alterations in pericentrin localization in cells lacking vimentin, we compared the distribution of Rab11 in wt and vimentin knockout cells. As it is shown in Fig. 7, MEF wt display a well defined Rab11 pattern, forming accumulations around the points of origin of cilia, marked by acetylated tubulin. In contrast, staining of MEF *Vim(-/-)* with anti-Rab11 revealed a disperse Rab11 distribution, with scarcer and less intense accumulations at the cilia origins, indicative of a disorganized pericentriolar material (Fig. 7A). To confirm these observations in a different cell type, we obtained A549 cells in which vimentin expression was knocked out by CRISPR-double nickase techniques (A549 *VIM*KO) (Suppl. Fig. 4). Importantly, A549 *VIM*KO cells also displayed marked defects in Rab11 distribution (Fig. 7B), characterized by a higher cytoplasmic staining and failure to concentrate in the pericentriolar material. Indeed, quantitation of these observations indicated that the proportion of cells displaying well defined pericentriolar material, identified by Rab11 staining, was significantly decreased in vimentin knockout cells, both MEF *Vim(-/-)* and A549 *VIM*KO, with respect to their wild type counterparts (Fig. 7A and B, graphs). Notably, A549 *VIM*KO cells also showed a marked defect in cilia formation, since staining with anti-acetylated tubulin yielded almost no detectable defined cilia structures (Fig. 7B).

**Figure 7.**
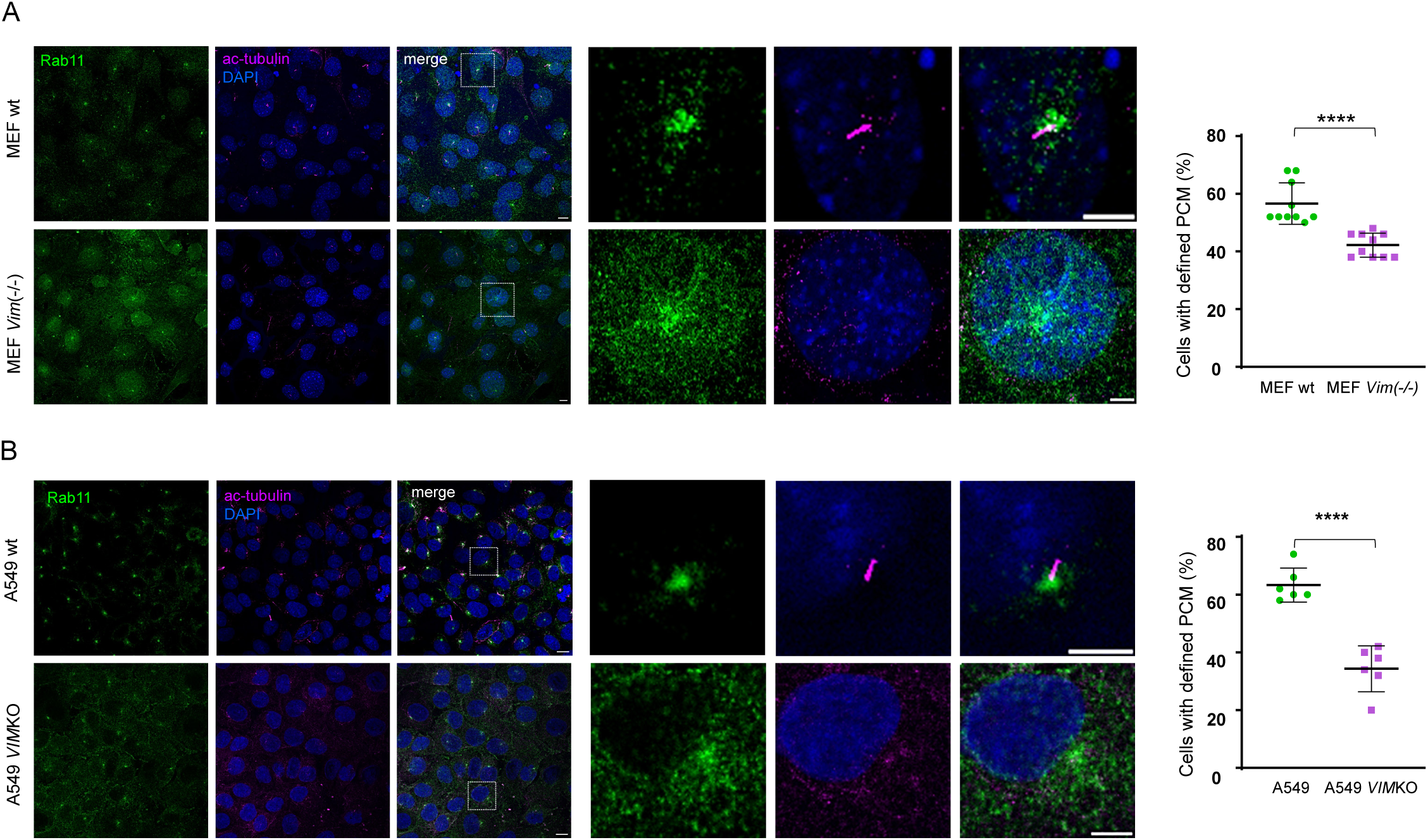
Impact of vimentin knockout on the distribution of Rab11. MEF wt or *Vim(-/-)* (A) or A549 wt and *VIM*KO cells (B) were stained for Rab11 and acetylated tubulin, as indicated. Single channels or merged images or overall projections are shown. The areas of interest marked by dotted boxes are enlarged in the right panels. The proportion of cells displaying a defined pericentriolar material (PCM) region, as highlighted by Rab11 staining are shown on the graphs at far right. Results shown are average values ± SEM of the proportions obtained in the analysis of ten (A) and six (B) fields, with approximately 50 cells each. ****p<0.0001 by unpaired *t*-test.

Taken together these results show that vimentin is required for the adequate distribution of Rab11 and other proteins involved in ciliogenesis.

## Discussion

The results shown herein unveil that abolishing vimentin expression leads to important defects in the morphology of the primary cilium, as well as to alterations in the levels or distribution of elements important for ciliogenesis, including γ-tubulin, pericentrin and Rab11. Some of the potential interactions of vimentin with elements involved in the formation of the primary cilium are schematized in Fig. 8. Together with the existing literature in this field, a hypothesis could be formulated, according to which the depletion of vimentin would provoke a traffic defect affecting the distribution of Rab11 vesicles and their targeting to the pericentriolar area. As Rab 11 is a critical regulator of ciliogenesis, its disruption could impair pericentrin localization and contribute to the altered pattern of γ-tubulin. In addition, the absence of vimentin can alter the pattern of acetylated tubulin. Therefore, vimentin depletion could affect primary cilia by disrupting both vesicular traffic and tubulin organization and modification, through independent or connected mechanisms, which require further investigation. In addition, effects on cell proliferation or cell cycle would need to be considered. In any event, as the primary cilium is a master sensing organelle which influences multiple cellular functions, these results unveil novel mechanisms for the complex effects of vimentin in cell biology.

**Figure 8.**
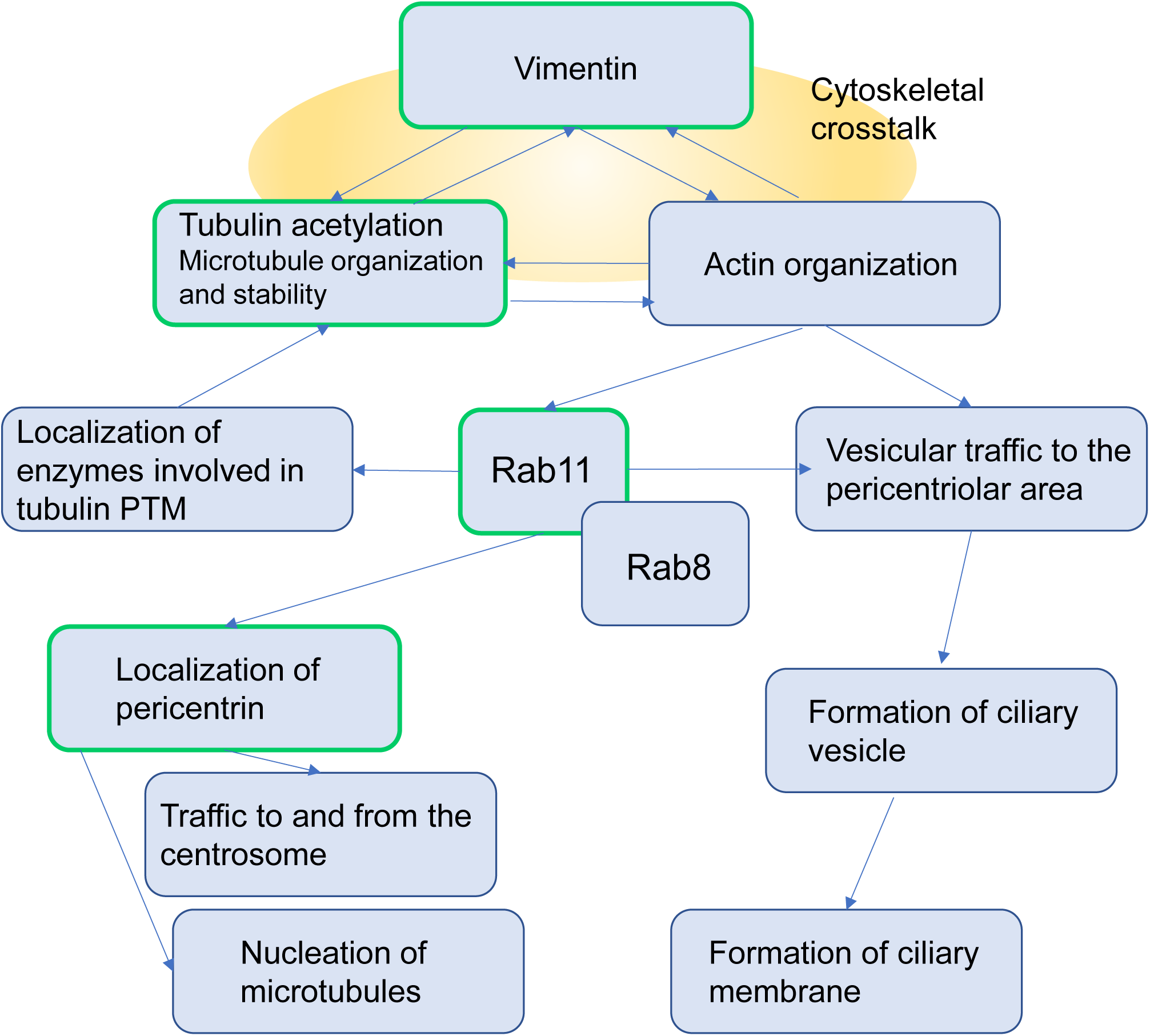
Potential interactions of vimentin with some of the elements involved in the formation of the primary cilium. The scheme depicts some of the elements or events reported in the literature to be involved in the formation of the primary cilium, which could be affected directly or indirectly by vimentin. See text for details. Of these, our observations indicate that depletion of vimentin affects the distribution of acetylated tubulin, γ-tubulin, Rab11 and pericentrin. PTM, posttranslational modifications.

We have observed that vimentin is present along the whole length of the primary cilium. Interestingly, our optical microscopy 3D reconstructions show a close intertwining of vimentin and acetylated tubulin throughout this structure. Remarkably, 3D reconstruction of electron microscopy data [16], had revealed the presence of filamentous densities along the cilium and in between the microtubules, which appear to stabilize the structure, and that to the best of our knowledge have not been unequivocally identified. Our present results suggest that vimentin could be part of these structures, probably together with other filamentous or cytoskeletal proteins, such as septins [29], and hypothetically could also play a role in cilium architecture. Vimentin localization and organization are strongly dependent on posttranslational modifications. Our results indicate that pSer56-vimentin is enriched in the cilium, and specially at the pericentriolar area of A549 cells. Vimentin Ser56 is a site for CDK1 phosphorylation during mitosis [38]. Indeed, we observe a widespread signal of pSer56-vimentin in mitotic cells. However, the localization of pSer56-vimentin at the primary cilium in interphase cells is a novel observation. Whether Ser56 phosphorylation targets vimentin to the primary cilium or is a consequence of its presence at this location, remains to be elucidated. Interestingly, during mitosis, vimentin phosphorylated at Ser56 by CDK1 recruits Plk1 and is further phosphorylated by this kinase at Ser82 [38]. Plk1 is an important kinase for the regulation of the cell cycle that also plays key roles in ciliary biology. In fact, both CDK1 and Plk1 have been involved in cilia disassembly [44, 45]. Moreover, vimentin is a substrate for Aurora A kinase, which is also involved in cilia resorption [22]. Therefore, as the primary cilium is a transient structure that is reabsorbed during mitosis, it would be interesting to ascertain whether there are posttranslational modifications, specially phosphorylation, that take place on vimentin in a cyclical manner and govern its presence and/or function at the primary cilium.

Tubulin posttranslational modifications are very important for ciliogenesis and affect cilium length [17, 46]. We have observed that the lack of vimentin also affects the distribution of acetylated tubulin and of γ-tubulin. Indeed, our results show that both forms appear in disorganized structures in vimentin knockout cells. Vimentin is involved in a bidirectional interplay with microtubules and microtubule modifications. Vimentin filaments template and stabilize microtubule networks [47], and in turn, microtubules transport intermediate filament precursors in the cell [48]. Nevertheless, evidence available offers some conflicting data, since vimentin has been reported both to increase and decrease tubulin acetylation, which could depend on the experimental context. Vimentin overexpression has been reported to decrease α-tubulin acetylation levels in HeLa cells [49]. On the other hand, vimentin appears to increase and stabilize acetylated microtubules in MEF [50]. In migrating astrocytes, vimentin filaments have been reported to dampen a positive feedback loop between microtubule acetylation and Rho-driven actomyosin contractility, [51]. A potential mechanism for this has been proposed involving an increase in tubulin acetylation upon vimentin depletion, the release of a Rho GEF from acetylated tubulin, and consequently, the activation of the Rho-RhoK pathway. This would contribute to a positive impact of vimentin depletion on actomyosin contractility, which has been observed in several vimentin deficient cellular models [52, 53]. In turn, microtubule acetylation has been reported to affect the transport and organization of intermediate filaments [48, 54]. Here, we have observed that vimentin depletion affects acetylated tubulin distribution in MEF and A549 cells. Nevertheless, further studies will be required to fully understand this complex interplay.

The vimentin monomer possesses a conserved single cysteine residue, C328 in human vimentin, that is important for the optimal performance of the protein in filament assembly and organelle positioning [33]. This residue is also a key target for oxidative modifications and subsequent disruption of the vimentin network in cells [2, 55]. In previous works, we have observed that cells expressing a vimentin C328S mutant show defects in MTOC/centrosomal organization [33], characterized by a less defined pattern of γ-tubulin compared to cells expressing vimentin wt. In addition, the cysteine deficient vimentin does not properly support the accumulation of γ-tubulin at aggresome-like structures upon inhibition of the proteasome, as does the wild type protein [33]. In view of the results presented herein, it would be interesting to assess whether assembly defective vimentin mutants can impact the formation of the primary cilium.

Interestingly, our results show that the levels of pericentrin are lower in MEF *Vim(-/-)*, as detected by immunofluorescence. This could also contribute to defective cilia formation. In fact, pericentrin depletion has been shown to disrupt cilia formation in human retinal epithelial (RPE1) cells [41]. Our observations are consistent with a recent report indicating that vimentin “mediates” the structure of the pericentriolar material and its absence correlates with a decrease of pericentrin at the centrosome [50]. On the other hand, elevated pericentrin, as observed in trisomy 21 delays primary ciliogenesis by disrupting multiple early steps of this process, and decreases sonic hedgehog signaling [40]. This indicates that pericentrin levels and distribution need to be tightly controlled for correct cilia formation.

Vesicular traffic plays a key role in the dynamics of the cilium by regulating the arrival and shuttling of building materials [56]. In particular, Rab11 and Rab8 GTPases have been shown to act in a concerted manner in ciliogenesis by modulating directional traffic and coupling cargo transport by recycling endosomes (Rab11) to subsequent vesicle docking at the plasma membrane (Rab8) [20]. Rab11 depletion through siRNA results in a decrease in cilia length, apparently through a reduced activation of Rab8, which has been reported to act downstream from Rab11 [20, 21]. Moreover, Rab11 has been shown to regulate the concentration of pericentrin at the centrosome [43], and participate in the traffic of certain enzymes involved in tubulin posttranslational modifications. In turn, association of Rab11 with the mother centriole appendages has been reported to influence its activity and therefore, the traffic of recycling endosomes at this location [57]. On the other hand, Rab7 is necessary for cilia disassembly [58]. Interestingly, Rab7 overexpression has been reported to interact with and regulate vimentin phosphorylation at Ser39 and 56 [59].

Vimentin is necessary for vesicular traffic. Vimentin deficient cells display an abnormal distribution of endolysosomes [33, 60, 61]. This is also the case in cells expressing certain vimentin mutants [33]. Nevertheless, the functional connection between vimentin and Rab11-mediated vesicular traffic has not been explored to the best of our knowledge. Our results show that the distribution of Rab11 is profoundly altered both in MEF *Vim(-/-)* and in A549 *VIM*KO cells. In the absence of vimentin, Rab11 structures appear disperse throughout the cytoplasm, and cells lack the defined accumulation of Rab11 in the pericentriolar area typical of cells expressing vimentin.

A potential link between vimentin and Rab11 mediated transport could involve the modulation of actomyosin contractility. As stated above, vimentin has been reported to exert a negative effect on actin polymerization in several experimental models, both under resting conditions as well as in response to stimulation with serum or with electrophilic compounds [52, 53]. Indeed, vimentin deficiency is accompanied by increased f-actin, both in interphase and in mitotic cells [52, 53, 62]. In mitosis, interference with the appropriate localization of vimentin profoundly alters the organization of the actin cortex and hampers normal cell division [62, 63]. Actin is a major regulator of both vesicular traffic and cilia formation [26, 64, 65]. Interestingly, a recent work has shown that impairment of actin polymerization at the apical region of the cell, either pharmacologically or upon caveolin-1 depletion, leads to increased cilia length, in association with an increase in the presence of Rab11 vesicles in the pericentriolar area [26]. The authors propose that f-actin clearing at the ciliary base facilitates the arrival of more transport vesicles, providing extra material for cilia growth [26]. Taken together, these observations suggest the possibility that the negative effect of knocking out vimentin on cilia formation, could be related, at least in part, to the modulation of actomyosin contractility.

Finally, the complexity of cytoskeletal crosstalk is being increasingly unveiled. Several actin remodeling proteins, as well as actomyosin-mediated mechanosensing can control microtubule acetylation and organization [51, 66, 67], further exposing the interdependence of these systems (Fig. 8). Accumulating evidence indicates that interfering with any of the major cytoskeletal networks will alter their delicate equilibrium, affecting essential cellular processes. The results reported herein indicate that the intermediate filament protein vimentin plays a significant role in the process of ciliogenesis, emerging as a novel element in the formation and/or architecture of the primary cilium, potentially participating in cytoskeletal crosstalk at this structure.

## Acknowledgements

We are indebted to the personnel from the Laser Confocal and Multidimensional in vivo Microscopy, Electronic Microscopy and Proteomics and Genomics facilities of CIB Margarita Salas, for expert assistance.

## References

[1] S. Etienne-Manneville, Cytoplasmic Intermediate Filaments in Cell Biology, Annu Rev Cell Dev Biol 34 (2018) 1–28.

[2] D. Pérez-Sala, R. Quinlan, The redox-responsive roles of intermediate filaments in cellular stress detection, integration and mitigation, Curr Opin Cell Biol 86 (2024) 102283.

[3] I. Ramos, K. Stamatakis, C.L. Oeste, D. Perez-Sala, Vimentin as a Multifaceted Player and Potential Therapeutic Target in Viral Infections, International journal of molecular sciences 21(13) (2020) 4675.

[4] A.E. Patteson, A. Vahabikashi, R.D. Goldman, P.A. Janmey, Mechanical and Non-Mechanical Functions of Filamentous and Non-Filamentous Vimentin, Bioessays 42(11) (2020) e2000078.

[5] G. Colakoglu, A. Brown, Intermediate filaments exchange subunits along their length and elongate by end-to-end annealing, J Cell Biol 185(5) (2009) 769–77.

[6] J.E. Eriksson, T. He, A.V. Trejo-Skalli, A.-S. Härmälä-Braskén, J. Hellman, Y.-H. Chou, R.D. Goldman, Specific in vivo phosphorylation sites determine the assembly dynamics of vimentin intermediate filaments, J Cell Sci 117 (2004) 919–932.

[7] N.T. Snider, M.B. Omary, Post-translational modifications of intermediate filament proteins: mechanisms and functions, Nat Rev Mol Cell Biol 15(3) (2014) 163–77.

[8] E. Griesser, V. Vemula, A. Mónico, D. Pérez-Sala, M. Fedorova, Dynamic posttranslational modifications of cytoskeletal proteins unveil hot spots under nitroxidative stress, Redox biology 44 (2021) 102014.

[9] L. Chang, R.D. Goldman, Intermediate filaments mediate cytoskeletal crosstalk, Nat Rev Mol Cell Biol 5(8) (2004) 601–13.

[10] F. Huber, A. Boire, M.P. Lopez, G.H. Koenderink, Cytoskeletal crosstalk: when three different personalities team up, Curr Opin Cell Biol 32 (2015) 39–47.

[11] A.B. Ndiaye, G.H. Koenderink, M. Shemesh, Intermediate Filaments in Cellular Mechanoresponsiveness: Mediating Cytoskeletal Crosstalk From Membrane to Nucleus and Back, Frontiers in cell and developmental biology 10 (2022) 882037.

[12] S. Parvanian, L.S. Coelho-Rato, A.E. Patteson, J.E. Eriksson, Vimentin takes a hike - Emerging roles of extracellular vimentin in cancer and wound healing, Curr Opin Cell Biol 85 (2023) 102246.

[13] L. Suprewicz, M. Swoger, S. Gupta, E. Piktel, F.J. Byfield, D.V. Iwamoto, D. Germann, J. Reszec, N. Marcinczyk, R.J. Carroll, P.A. Janmey, J.M. Schwarz, R. Bucki, A.E. Patteson, Extracellular Vimentin as a Target Against SARS-CoV-2 Host Cell Invasion, Small 18(6) (2022) e2105640.

[14] V. Lalioti, S. González-Sanz, I. Lois-Bermejo, P. González-Jiménez, A. Viedma-Poyatos, A. Merino, M.A. Pajares, D. Pérez-Sala, Cell surface detection of vimentin, ACE2 and SARS-CoV-2 Spike proteins reveals selective colocalization at primary cilia, Scientific reports 12 (2022) 7063.

[15] K.I. Hilgendorf, C.T. Johnson, P.K. Jackson, The primary cilium as a cellular receiver: organizing ciliary GPCR signaling, Curr Opin Cell Biol 39 (2016) 84–92.

[16] S. Sun, R.L. Fisher, S.S. Bowser, B.T. Pentecost, H. Sui, Three-dimensional architecture of epithelial primary cilia, Proc Natl Acad Sci U S A 116(19) (2019) 9370–9379.

[17] K. He, K. Ling, J. Hu, The emerging role of tubulin posttranslational modifications in cilia and ciliopathies, Biophys Rep 6 (2020) 89–104.

[18] L. Labat-de-Hoz, A. Rubio-Ramos, J. Casares-Arias, M. Bernabe-Rubio, I. Correas, M.A. Alonso, A Model for Primary Cilium Biogenesis by Polarized Epithelial Cells: Role of the Midbody Remnant and Associated Specialized Membranes, Frontiers in cell and developmental biology 8 (2020) 622918.

[19] D.K. Breslow, A.J. Holland, Mechanism and Regulation of Centriole and Cilium Biogenesis, Annu Rev Biochem 88 (2019) 691–724.

[20] A. Knodler, S. Feng, J. Zhang, X. Zhang, A. Das, J. Peranen, W. Guo, Coordination of Rab8 and Rab11 in primary ciliogenesis, Proc Natl Acad Sci U S A 107(14) (2010) 6346–51.

[21] C.J. Westlake, L.M. Baye, M.V. Nachury, K.J. Wright, K.E. Ervin, L. Phu, C. Chalouni, J.S. Beck, D.S. Kirkpatrick, D.C. Slusarski, V.C. Sheffield, R.H. Scheller, P.K. Jackson, Primary cilia membrane assembly is initiated by Rab11 and transport protein particle II (TRAPPII) complex-dependent trafficking of Rabin8 to the centrosome, Proc Natl Acad Sci U S A 108(7) (2011) 2759–64.

[22] G. Habeck, J. Schweiggert, Proteolytic control in ciliogenesis: Temporal restriction or early initiation?, Bioessays 44(9) (2022) e2200087.

[23] J.J. Moser, M.J. Fritzler, Y. Ou, J.B. Rattner, The PCM-basal body/primary cilium coalition, Seminars in cell & developmental biology 21(2) (2010) 148–55.

[24] A. Vertii, H. Hehnly, S. Doxsey, The Centrosome, a Multitalented Renaissance Organelle, Cold Spring Harbor perspectives in biology 8(12) (2016) a025049.

[25] N. Schweizer, J. Luders, From tip to toe - dressing centrioles in gammaTuRC, J Cell Sci 134(14) (2021) jcs258397.

[26] L. Rangel, M. Bernabe-Rubio, J. Fernandez-Barrera, J. Casares-Arias, J. Millan, M.A. Alonso, I. Correas, Caveolin-1alpha regulates primary cilium length by controlling RhoA GTPase activity, Scientific reports 9(1) (2019) 1116.

[27] C.E.L. Smith, A.V.R. Lake, C.A. Johnson, Primary Cilia, Ciliogenesis and the Actin Cytoskeleton: A Little Less Resorption, A Little More Actin Please, Frontiers in cell and developmental biology 8 (2020) 622822.

[28] Q. Hu, L. Milenkovic, H. Jin, M.P. Scott, M.V. Nachury, E.T. Spiliotis, W.J. Nelson, A septin diffusion barrier at the base of the primary cilium maintains ciliary membrane protein distribution, Science 329(5990) (2010) 436–9.

[29] M.S. Kim, C.D. Froese, H. Xie, W.S. Trimble, Immunofluorescent staining of septins in primary cilia, Methods in cell biology 136 (2016) 269–83.

[30] Y. Nishimura, K. Kasahara, M. Inagaki, Intermediate filaments and IF-associated proteins: from cell architecture to cell proliferation, Proceedings of the Japan Academy. Series B, Physical and biological sciences 95(8) (2019) 479–493.

[31] K. Tateishi, T. Nishida, K. Inoue, S. Tsukita, Three-dimensional Organization of Layered Apical Cytoskeletal Networks Associated with Mouse Airway Tissue Development, Scientific reports 7 (2017) 43783.

[32] K. Stamatakis, F.J. Sánchez-Gómez, D. Pérez-Sala, Identification of novel protein targets for modification by 15-deoxy-D12,14-prostaglandin J2 in mesangial cells reveals multiple interactions with the cytoskeleton., J Am Soc Nephrol 17 (2006) 89–98.

[33] D. Pérez-Sala, C.L. Oeste, A.E. Martínez, B. Garzón, M.J. Carrasco, F.J. Cañada, Vimentin filament organization and stress sensing depend on its single cysteine residue and zinc binding, Nature communications 6 (2015) 7287.

[34] K.A. Mitchell, Isolation of primary cilia by shear force, Current protocols in cell biology Chapter 3 (2013) Unit 3 42 1–9.

[35] I. Cristobo, M.J. Larriba, V. de los Rios, F. Garcia, A. Munoz, J.I. Casal, Proteomic analysis of 1alpha,25-dihydroxyvitamin D3 action on human colon cancer cells reveals a link to splicing regulation, J Proteomics 75(2) (2011) 384–97.

[36] B.T. Helfand, M.G. Mendez, S.N. Murthy, D.K. Shumaker, B. Grin, S. Mahammad, U. Aebi, T. Wedig, Y.I. Wu, K.M. Hahn, M. Inagaki, H. Herrmann, R.D. Goldman, Vimentin organization modulates the formation of lamellipodia, Mol Biol Cell 22(8) (2011) 1274–89.

[37] H. Goto, H. Kosako, K. Tanabe, M. Yanagida, M. Sakurai, M. Amano, K. Kaibuchi, M. Inagaki, Phosphorylation of vimentin by Rho-associated kinase at a unique amino-terminal site that is specifically phosphorylated during cytokinesis, J Biol Chem 273(19) (1998) 11728–36.

[38] T. Yamaguchi, H. Goto, T. Yokoyama, H. Sillje, A. Hanisch, A. Uldschmid, Y. Takai, T. Oguri, E.A. Nigg, M. Inagaki, Phosphorylation by Cdk1 induces Plk1-mediated vimentin phosphorylation during mitosis, J Cell Biol 171(3) (2005) 431–6.

[39] S. Shankar, Z.T. Hsu, A. Ezquerra, C.C. Li, T.L. Huang, E. Coyaud, R. Viais, C. Grauffel, B. Raught, C. Lim, J. Luders, S.Y. Tsai, K.C. Hsia, Alpha gamma-tubulin complex-dependent pathway suppresses ciliogenesis by promoting cilia disassembly, Cell reports 41(7) (2022) 111642.

[40] C.E. Jewett, B.L. McCurdy, E.T. O’Toole, A.J. Stemm-Wolf, K.S. Given, C.H. Lin, V. Olsen, W. Martin, L. Reinholdt, J.M. Espinosa, K.D. Sullivan, W.B. Macklin, R. Prekeris, C.G. Pearson, Trisomy 21 induces pericentriolar crowding delaying primary ciliogenesis and mouse cerebellar development, eLife 12 (2023) e78202.

[41] A. Jurczyk, A. Gromley, S. Redick, J. San Agustin, G. Witman, G.J. Pazour, D.J. Peters, S. Doxsey, Pericentrin forms a complex with intraflagellar transport proteins and polycystin-2 and is required for primary cilia assembly, J Cell Biol 166(5) (2004) 637–43.

[42] H. Hehnly, S. Doxsey, Rab11 endosomes contribute to mitotic spindle organization and orientation, Dev Cell 28(5) (2014) 497–507.

[43] N. Krishnan, M. Swoger, L.I. Rathbun, P.J. Fioramonti, J. Freshour, M. Bates, A.E. Patteson, H. Hehnly, Rab11 endosomes and Pericentrin coordinate centrosome movement during pre-abscission in vivo, Life science alliance 5(7) (2022) e202201362.

[44] G. Wang, Q. Chen, X. Zhang, B. Zhang, X. Zhuo, J. Liu, Q. Jiang, C. Zhang, PCM1 recruits Plk1 to the pericentriolar matrix to promote primary cilia disassembly before mitotic entry, J Cell Sci 126(Pt 6) (2013) 1355–65.

[45] L. Wang, B.D. Dynlacht, The regulation of cilium assembly and disassembly in development and disease, Development 145(18) (2018) dev151407.

[46] P. Guichard, M.H. Laporte, V. Hamel, The centriolar tubulin code, Seminars in cell & developmental biology 137 (2023) 16–25.

[47] Z. Gan, L. Ding, C.J. Burckhardt, J. Lowery, A. Zaritsky, K. Sitterley, A. Mota, N. Costigliola, C.G. Starker, D.F. Voytas, J. Tytell, R.D. Goldman, G. Danuser, Vimentin Intermediate Filaments Template Microtubule Networks to Enhance Persistence in Cell Polarity and Directed Migration, Cell systems 3(3) (2016) 252–263 e8.

[48] C. Hookway, L. Ding, M.W. Davidson, J.Z. Rappoport, G. Danuser, V.I. Gelfand, Microtubule-dependent transport and dynamics of vimentin intermediate filaments, Mol Biol Cell 26(9) (2015) 1675–86.

[49] P. Liu, S. Zhang, J. Ma, D. Jin, Y. Qin, M. Chen, Vimentin inhibits alpha-tubulin acetylation via enhancing alpha-TAT1 degradation to suppress the replication of human parainfluenza virus type 3, PLoS pathogens 18(9) (2022) e1010856.

[50] R. Saldanha, M. Tri Ho Thanh, N. Krishnan, H. Hehnly, A.E. Patteson, Vimentin supports cell polarization by enhancing centrosome function and microtubule acetylation, bioRxiv (2023) doi: 10.1101/2023.02.17.528977.

[51] S. Seetharaman, B. Vianay, V. Roca, A.J. Farrugia, C. De Pascalis, B. Boeda, F. Dingli, D. Loew, S. Vassilopoulos, A. Bershadsky, M. Thery, S. Etienne-Manneville, Microtubules tune mechanosensitive cell responses, Nat Mater 21(3) (2022) 366–377.

[52] Y. Jiu, J. Peranen, N. Schaible, F. Cheng, J.E. Eriksson, R. Krishnan, P. Lappalainen, Vimentin intermediate filaments control actin stress fiber assembly through GEF-H1 and RhoA, J Cell Sci 130(5) (2017) 892–902.

[53] P. González-Jiménez, S. Duarte, A. Martínez-Fernández, E. Navarro-Carrasco, V. Lalioti, M.A. Pajares, D. Pérez-Sala, Vimentin single cysteine residue acts as a tunable sensor for network organization and as a key for actin remodeling in response to oxidants and electrophiles, Redox biology 64 (2023) 102756.

[54] L.S. Rathje, N. Nordgren, T. Pettersson, D. Ronnlund, J. Widengren, P. Aspenstrom, A.K. Gad, Oncogenes induce a vimentin filament collapse mediated by HDAC6 that is linked to cell stiffness, Proc Natl Acad Sci U S A 111(4) (2014) 1515–20.

[55] A. Mónico, S. Duarte, M.A. Pajares, D. Pérez-Sala, Vimentin disruption by lipoxidation and electrophiles: role of the cysteine residue and filament dynamics, Redox biology 23 (2019) 101098.

[56] S. Yoshimura, J. Egerer, E. Fuchs, A.K. Haas, F.A. Barr, Functional dissection of Rab GTPases involved in primary cilium formation, J Cell Biol 178(3) (2007) 363–9.

[57] H. Hehnly, C.T. Chen, C.M. Powers, H.L. Liu, S. Doxsey, The centrosome regulates the Rab11-dependent recycling endosome pathway at appendages of the mother centriole, Curr Biol 22(20) (2012) 1944–50.

[58] G. Wang, H.B. Hu, Y. Chang, Y. Huang, Z.Q. Song, S.B. Zhou, L. Chen, Y.C. Zhang, M. Wu, H.Q. Tu, J.F. Yuan, N. Wang, X. Pan, A.L. Li, T. Zhou, X.M. Zhang, K. He, H.Y. Li, Rab7 regulates primary cilia disassembly through cilia excision, J Cell Biol 218(12) (2019) 4030–4041.

[59] L. Cogli, C. Progida, R. Bramato, C. Bucci, Vimentin phosphorylation and assembly are regulated by the small GTPase Rab7a, Biochim Biophys Acta 1833(6) (2013) 1283–93.

[60] M.L. Styers, G. Salazar, R. Love, A.A. Peden, A.P. Kowalczyk, V. Faundez, The endo-lysosomal sorting machinery interacts with the intermediate filament cytoskeleton, Mol Biol Cell 15(12) (2004) 5369–82.

[61] C.L. Oeste, M. Martínez-López, D. Pérez-Sala, Taking a lipidation-dependent path towards endolysosomes, Commun Integr Biol 8(5) (2015) e1078041.

[62] S. Duarte, A. Viedma-Poyatos, E. Navarro-Carrasco, A.E. Martinez, M.A. Pajares, D. Perez-Sala, Vimentin filaments interact with the actin cortex in mitosis allowing normal cell division, Nature communications 10 (2019) 4200.

[63] M.P. Serres, M. Samwer, B.A. Truong Quang, G. Lavoie, U. Perera, D. Gorlich, G. Charras, M. Petronczki, P.P. Roux, E.K. Paluch, F-Actin Interactome Reveals Vimentin as a Key Regulator of Actin Organization and Cell Mechanics in Mitosis, Dev Cell 52(2) (2020) 210–222 e7.

[64] A.J. Ridley, Rho GTPases and actin dynamics in membrane protrusions and vesicle trafficking, Trends Cell Biol 16(10) (2006) 522–9.

[65] M.L. Drummond, M. Li, E. Tarapore, T.T.L. Nguyen, B.J. Barouni, S. Cruz, K.C. Tan, A.E. Oro, S.X. Atwood, Actin polymerization controls cilia-mediated signaling, J Cell Biol 217(9) (2018) 3255–3266.

[66] J. Fernandez-Barrera, M.A. Alonso, Coordination of microtubule acetylation and the actin cytoskeleton by formins, Cell Mol Life Sci 75(17) (2018) 3181–3191.

[67] P. Shi, Y. Wang, Y. Huang, C. Zhang, Y. Li, Y. Liu, T. Li, W. Wang, X. Liang, C. Wu, Arp2/3-branched actin regulates microtubule acetylation levels and affects mitochondrial distribution, J Cell Sci 132(6) (2019) jcs226506.

