## Supplementary material for "Key role of vimentin in the organization of the primary cilium": Suppl. Fig.

Running title: Vimentin at the primary cilium

### Supplementary Figures

Supplementary Information

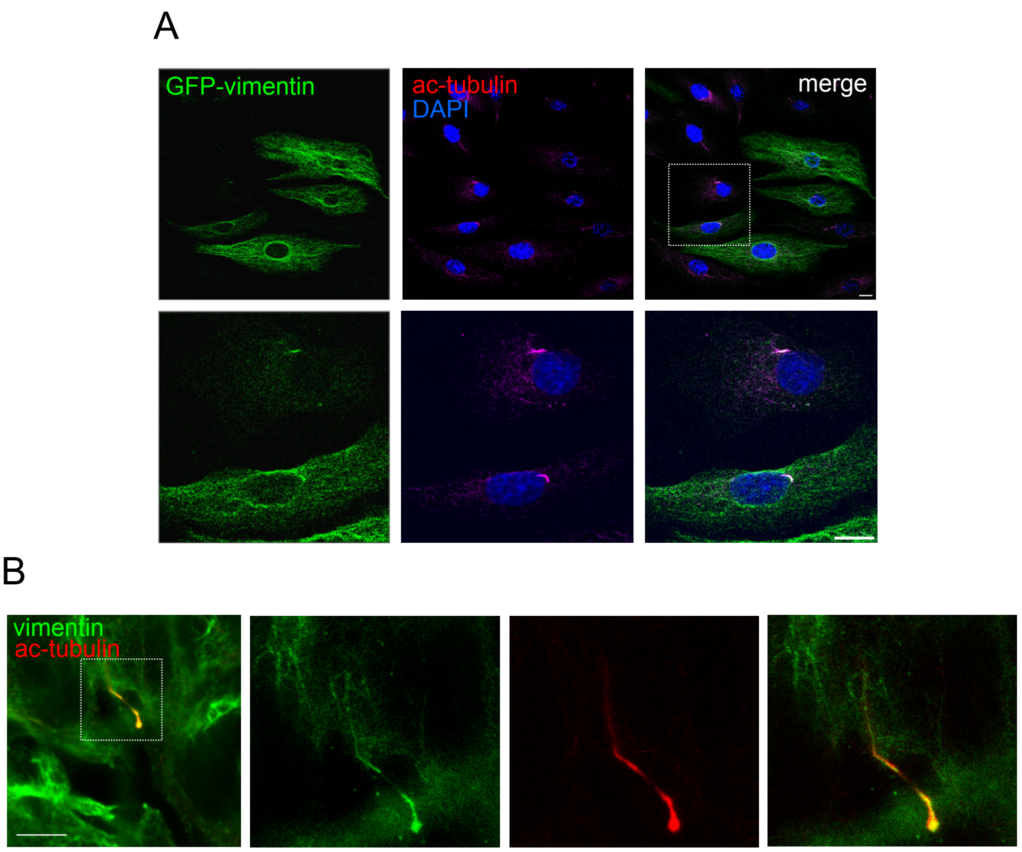


**Supplementary figure 1. Presence of vimentin at the primary cilium.** (A) A549 cells were transfected with GFP-vimentin and acetylated tubulin was visualized by immunofluorescence. Nuclei were counterstained with DAPI. Images shown are middle stacks of single channels or merged images, as indicated. The region of interest delimited by the dotted square is enlarged in the images in the bottom row (scale bar, 10 µm). (B) Analysis of the presence of endogenous vimentin at the primary cilium of A549 cells by immunofluorescence and STED superresolution microscopy. Here, the projection of three sections is used to show a longer extension of the cilium length. Scale bar, 5 µm.

Supplementary Information

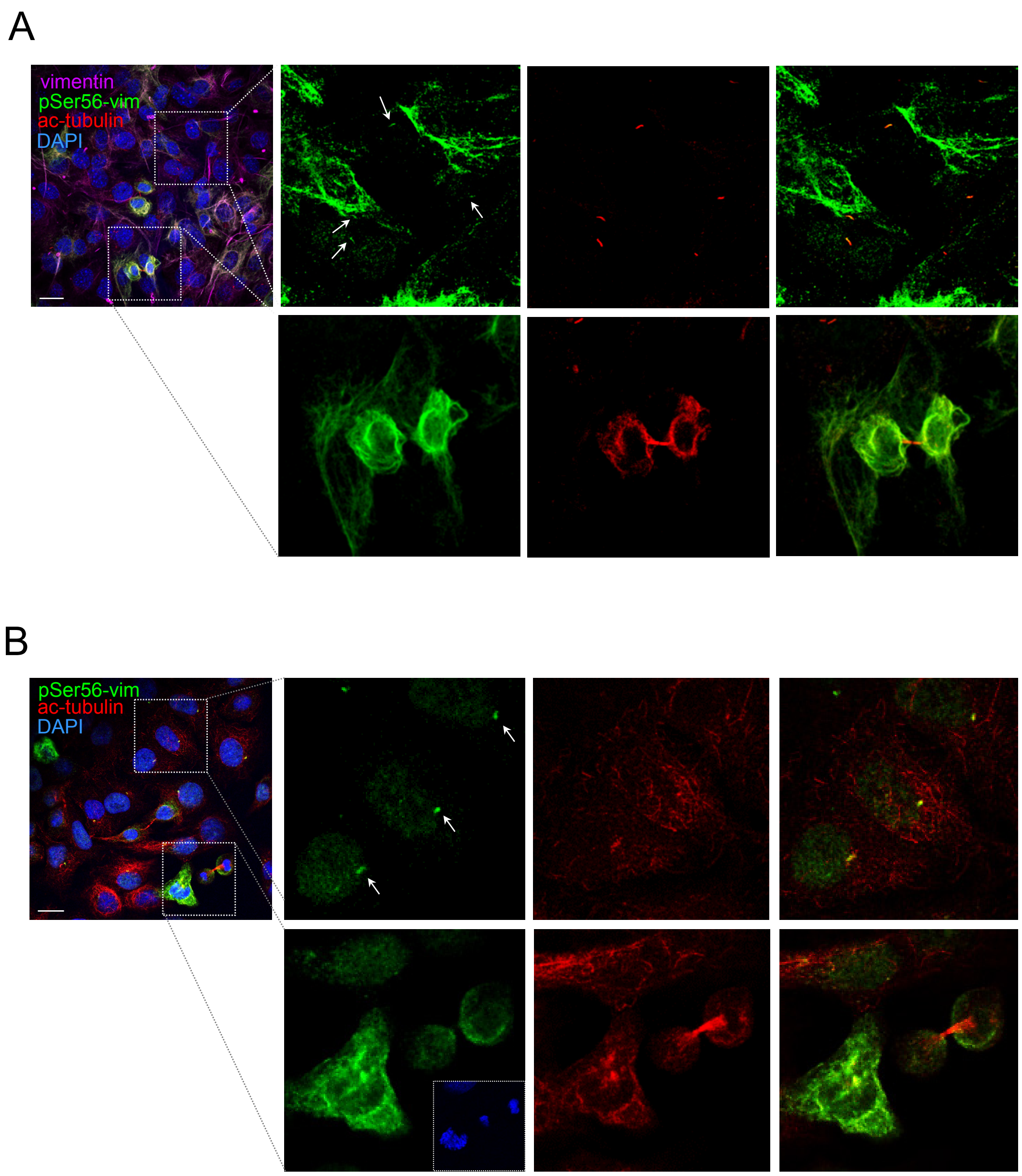


**Supplementary figure 2. Presence of pSer56-vimentin at various structures in MEF wt and A549 cells.** (A) The presence of total vimentin, detected with the E5 antibody, pSer56-vimentin, and acetylated tubulin, was assessed in MEF wt by immunofluorescence. (B) A549 cells were processed for the detection of pSer56-vimentin and acetylated tubulin. In (A) and (B) nuclei were counterstained with DAPI. The images on the left show overlays of all channels. The areas or interest indicated by dotted boxes are enlarged at the right to illustrate the coincidence of the pSer56-vimentin and acetylated tubulin signals at primary cilia (arrows), or at the bottom, to illustrate the presence of pSer56-vimentin in cells at various stages of mitosis. All images are overall projections except those shown in (A), right images, upper row. In (A) upper images, the signal of pSer56-vimentin is deliberately overexposed in order to appreciate its presence at the primary cilium. In (B) lower images, an inset with DAPI staining is provided to identify the mitotic cells. Scale bars, 20 µm.

Supplementary Information

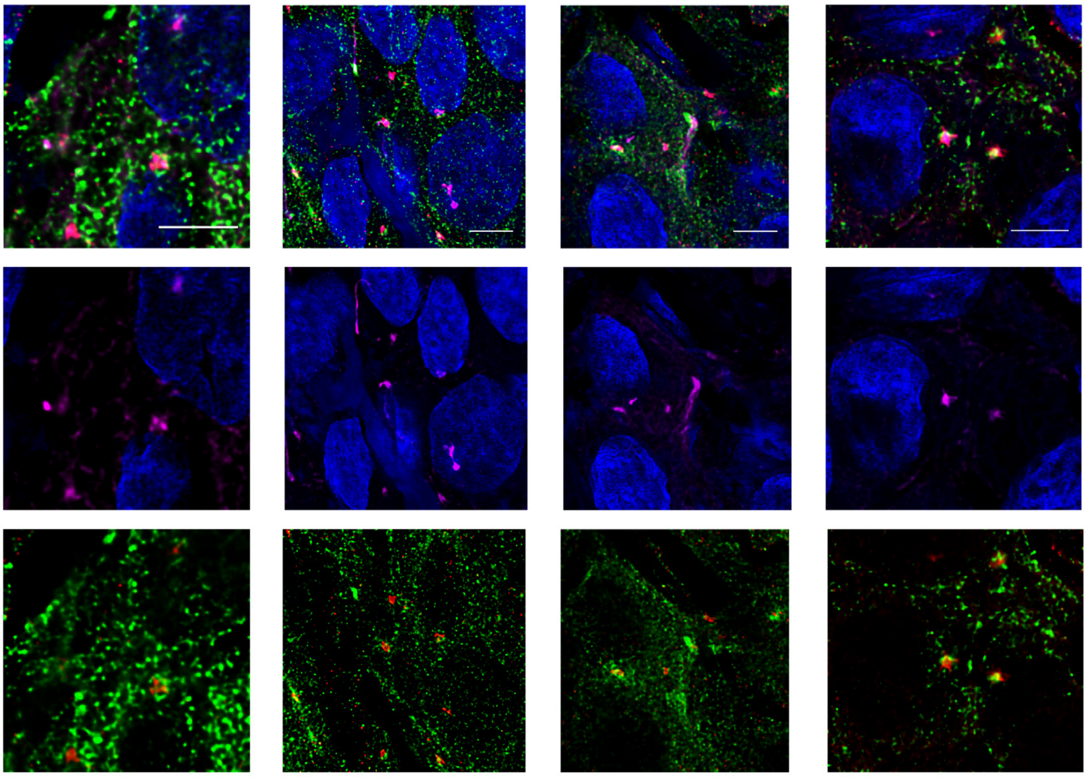

**Supplementary figure 3. Detection of pSer56-vimentin at the pericentriolar area in A549 cells.** Cilia were visualized by staining with anti-acetylated tubulin (ac-tubulin) and the pericentriolar material by Rab11 staining. Nuclei are counterstained with DAPI. Note the presence of pSer56-vimentin at areas enriched or encircled by the Rab11 signal. Images shown are overlays of single sections. Scale bars, 5 µm.

Supplementary Information

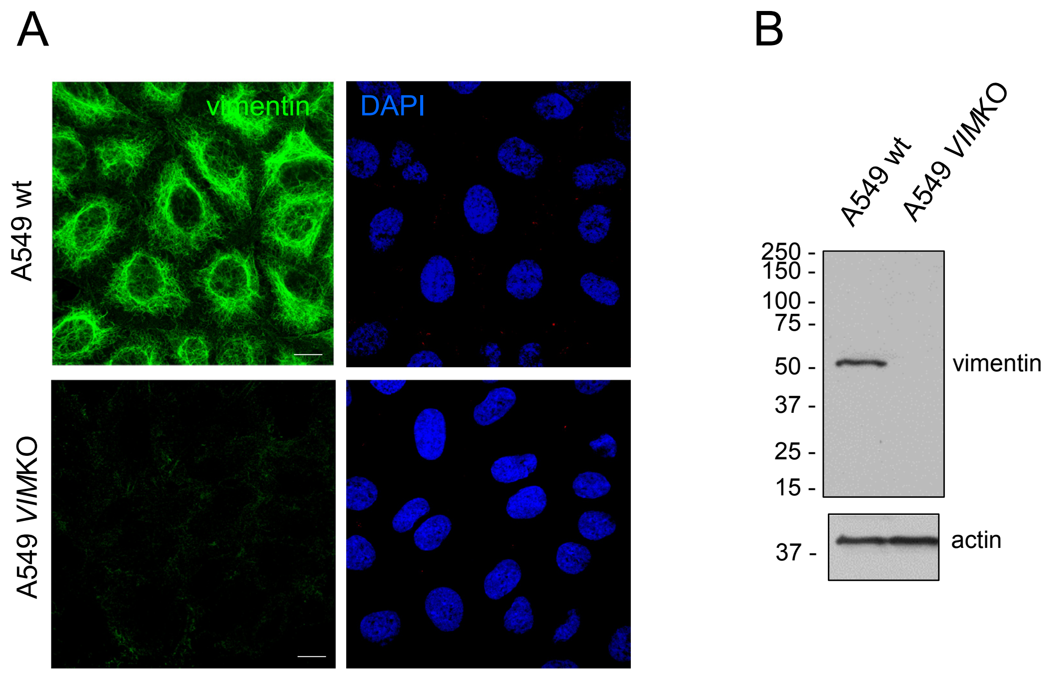


**Supplementary figure 4. Characterization of A549 *VIM*KO cells.** (A) Total vimentin was detected by immunofluorescence with the 84.1 antibody in A549 wt or *VIM*KO cells. Nuclei were counterstained with DAPI. Images are overall projections. Scale bars, 10 µm. (B) Total cell lysates from A549 wt or *VIM*KO cells containing 30 µg of protein were resolved by SDS-PAGE and levels of vimentin were assessed by western blot with the V9 antibody. Levels of actin are shown in the lower panel as a control. The position of molecular weight markers (in kDa) is shown at the left of blots.
